# Biotransformation of 2,4-dinitrotoluene in a phototrophic co-culture of engineered *Synechococcus elongatus* and *Pseudomonas putida*

**DOI:** 10.1101/404988

**Authors:** Derek T. Fedeson, Pia Saake, Patricia Calero, Pablo Iván Nikel, Daniel C. Ducat

## Abstract

In contrast to the current paradigm of using microbial monocultures in most biotechnological applications, increasing efforts are being directed towards engineering mixed-species consortia to perform functions that are difficult to program into individual strains. Additionally, the division of labor between specialist species found in natural consortia can lead to increased catalytic efficiency and stability relative to a monoculture or a community composed of generalists. In this work, we have designed a synthetic co-culture for phototrophic degradation of xenobiotics, composed of a cyanobacterium, (*Synechococcus elongatus* PCC 7942) and a heterotrophic bacterium (*Pseudomonas putida* EM173). Cyanobacteria fix CO_2_ through photosynthetic metabolism and secrete sufficient carbohydrates to support the growth and active metabolism of *P. putida*, which has been engineered to consume sucrose as the only carbon source and to degrade the environmental pollutant 2,4-dinitrotoluene (2,4-DNT). The synthetic consortium is able to degrade 2,4-DNT with only light and CO_2_ as inputs for the system, and it was stable over time through repeated backdilutions. Furthermore, cycling this consortium through low nitrogen medium promoted the accumulation of polyhydroxyalkanoate (PHA)–an added-value biopolymer–in *P. putida*, thus highlighting the versatility of this production platform. Altogether, the synthetic consortium allows for simultaneous bioproduction of PHA and remediation of the industrial pollutant 2,4-DNT, using light and CO_2_ as inputs.

**Importance:** In this study, we have created an artificial consortium composed of two bacterial species that enables the degradation of the industrially-produced environmental pollutant 2,4-DNT while simultaneously producing PHA bioplastic. In these co-cultures, the photosynthetic cyanobacteria fuel an engineered *P. putida* strain programmed both to use sucrose as a carbon source and to perform the biotransformation of 2,4-DNT. The division of labor in this synthetic co-culture is reminiscent of that commonly observed in microbial communities and represents a proof-of-principle example of how artificial consortia can be employed for bioremediation purposes. Furthermore, this co-culture system enabled the utilization of freshwater sources that could not be utilized in classical agriculture settings, reducing the potential competition of this alternative method of bioproduction with current agricultural practices, as well as remediation of contaminated water streams.

## Introduction

In nature, bacteria typically co-exist in communities with hundreds to thousands of other microorganisms, creating a complex web of inter-species metabolic reactions (1–4). Most consortia exhibit a high degree of “division of labor,” where individual species have specialized metabolisms and exchange metabolites and signals with neighbors (5, 6). Interactions range from the cooperative degradation of toluene (7, 8) to the consumption of metabolic waste products (9). Compartmentalization of metabolism across distinct species can confer metabolic capabilities on a consortium that may be difficult to engineer within any one individual. Additionally, natural microbial consortia frequently exhibit a high degree of robustness in the face of dynamic environmental conditions and are resilient to invasive microbes (10, 11). Thus, there has been increasing interest in rationally engineering microbial consortia for desired outputs by dividing metabolic pathways across species (12).

Previous work from our laboratory and others has focused on the utility of strains of cyanobacteria engineered to export photosynthetically-generated sucrose through the heterologous expression of the *cscB* gene from *Escherichia coli*, encoding sucrose permease (13–15). In one study utilizing this engineered strain of *Synechococcus elongatus* PCC7942 (hereafter, *S. elongatus* CscB), sucrose secretion was such that hypothetical scaled production would significantly exceed current productivities of traditional sugar crops like sugar cane, sugar beet, and corn (13). As such, the strain has been utilized on numerous occasions as a photosynthetic module in synthetic co-cultures as a supplier of fixed carbon for heterotrophic partners (16–20), including a recent report where co-cultures were maintained for longer than 5 months of continuous co-culture (21). Communally, these works have demonstrated that the *S. elongatus* CscB strain can be flexibly paired with a variety of heterotrophic bacteria [*Escherichia coli* W (16), *Bacillus subtilis* (16), *Azotobacter vinelandii* (18, 19), *Halomonas boliviensis* (21), and *Pseudomonas putida* (17)] and yeasts [*Saccharomyces cerevisiae* (13, 16), *Cryptococcus curvatus* (20), and *Rhodotorula glutinis* (20)] and demonstrated that these co-cultures can be used to photosynthetically produce valued biological products, *e.g.*, α-amylase (16), fatty acids (20), and polyhydroxyalkanoates (PHA) [*e.g.*, poly(3-hydroxybutyrate) (PHB) (17–19, 21)]. In work from Smith and Francis (19) as well as our lab (21) it was shown that *S. elongatus* CscB could be immobilized within hydrogels; this both enhanced carbon flux into sucrose production and allowed the cyanobacteria to exchange diffusible metabolites in medium, while enabling the heterotrophic cells to be readily harvested separately. Taken together, this co-culture approach has potential as a platform to enable the modular photosynthetic production of a flexible array of bioproducts.

A primary concern when considering large-scale aquatic based bioproduction, is that global potable water supplies are coming under increasing strain; scaled applications of algae or cyanobacteria would be more sustainable if they can utilize marginal waters not suitable for other agricultural purposes (22). Industrial wastewater streams are one such example, as they are often contaminated with chemical compounds toxic to both flora and fauna. 2,4-Dinitrotoluene (2,4-DNT) is a nitroaromatic compound and is one of a diverse array of xenobiotics that have been released into the environment due to industrial synthetic chemistry and other manufacturing processes (23). While 2,4-DNT is produced as a by-product during polyurethane and pesticide synthesis, one of the most significant sources of these contaminants is explosive manufacturing (24). One 2,4,6-trinitrotoluene (TNT)-manufacturing plant can contaminate five hundred thousand gallons of water with TNT and other nitroaromatics in a single day (25), and at some munitions manufacturing and processing sites, soil contamination is as high as 200 g of TNT per kilogram of soil (26). These compounds are highly stable within the environment, and remediation costs via incineration are estimated near $400 USD per cubic yard of soil (27).

Despite the fact that these nitroaromatic compounds have only recently been introduced into the environment, a number of bacterial strains capable of mineralizing these unusual chemical structures have been isolated (23, 24, 28). The most common mechanism by which 2,4-DNT is processed is through one or two-electron reduction via non-specific nitroreductases (29). Enzymes containing a redox-active flavin or iron center are especially prone to perform these reactions (30). Single electron reductions temporarily reduces the nitro group but it is then immediately re-oxidized in the presence of molecular oxygen resulting in the generation of a superoxide, the accumulation of which can lead to the ROS stress response (31). Two electron reductions result in the production of a toxic hydroxylamino derivative (Figure 3B) that can react with DNA and cause subsequent mutations (30). The nitroso- and hydroxylamino-derivatives are even more toxic than their parent molecule, able to form DNA and protein adducts that lead to mutagenesis and cellular damage (24, 32). Additionally, the derivatives from reduced nitroaromatics continue to persist in the environment (33), further emphasizing the need for alternative methods of degradation that allow for the complete biotransformation of these compounds.

**Figure 3:**
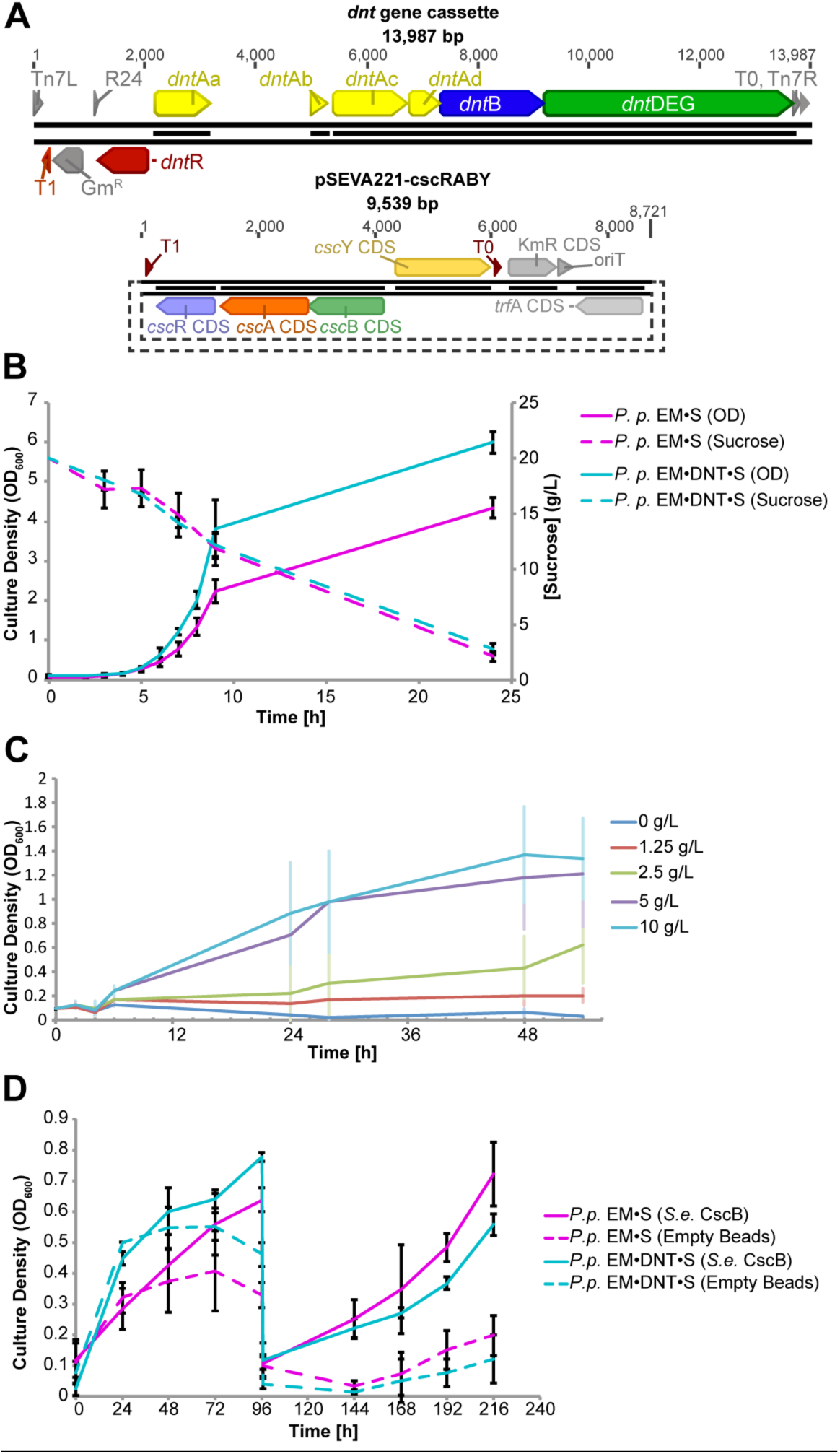
Growth characterization of sucrose-consuming *P. putida* EM·DNT·S strains. (A). Schematic representations of the 2,4-DNT degradation gene cluster from *Burkholderia* sp. R34 (top) and the gene cassette for sucrose utilization in pSEVA221-cscRABY (bottom). Growth of kinetics and sucrose utilization of the *P. putida* strains in 20 g/L of sucrose in M9 minimal medium. *n* = 4; error bars represent standard deviations. (C) Growth of *P. putida* EM·DNT·S at different concentrations of sucrose (0, 1.25, 2.5, 5, and 10 g/L) in M3 medium. (D) *P. putida* strains EM·S and *P. putida* EM·DNT·S were grown in M3 media with 20 g/L sucrose overnight and then inoculated at OD_600_ ∼ 0.1 into flasks containing alginate beads with or without encapsulated *S. elongatus* CscB, all cultures contained 1 mM IPTG for induction of sucrose export. OD_600_ measurements were taken over the course of 216 hours, tracking the growth of the *P. putida* strains in the co-culture. At 96 h post-inoculation, all of the M3 media was removed and replaced with fresh medium, allowing the residual *P. putida* cells on the surface of the alginate beads to repopulate the culture. For the experiments shown in (C) and (D), the mean values for *n* = 3 are indicated, and error bars represent standard deviations.

A separate, oxidative pathway for degrading 2,4-DNT was identified in *Burkholderia* sp. R34, a strain isolated from surface water contaminated by an ammunition waste plant (34). The gene cluster responsible for processing 2,4-DNT has been identified (34–36) and contains 7 genes. Enzymes within the *dnt* degradation pathway share homology with enzymes that function as part of an existing naphthalene degradation pathway in *Burkholderia sp*. R34 and proceed through oxidative steps that release nitrite. This suite of genes has since been chromosomally integrated into a strain of *Pseudomonas putida* EM173 via a Tn*7* construct to facilitate the analysis of this pathway (37, 38). *P. putida* is generally considered a reliable *chassis* for studying the biodegradation of organic compounds due to its tolerance of organic solvents (39) and versatile central metabolism (40, 41).

In this study, we explore whether the aforementioned synthetic consortium method (21) can be engineered to utilize and remediate water streams contaminated with the environmental pollutant 2,4-DNT while also producing the bioplastic polyhydroxyalkanoate (PHA) (Figure 1). This was accomplished through the pairing of an engineered strain of *P. putida*, containing the genes needed to both metabolize sucrose and to degrade 2,4-DNT with the sucrose-exporting *S. elongatus* CscB encapsulated in alginate hydrogel beads. Approaching the bioremediation of this compound via the synthetic consortium method allows for the bioprocess to be photosynthetically powered while avoiding the complexities of introducing a new enzymatic pathway into photosynthetic cyanobacteria. We demonstrated that these co-cultures can successfully execute the biotransformation of 2,4-DNT via the engineered pathway and are also able to accumulate PHA as a secondary function. This work is proof of principle for the use of synthetic cyanobacteria/heterotroph consortia for combined bioremediation and bioproduction applications.

**Figure 1:**
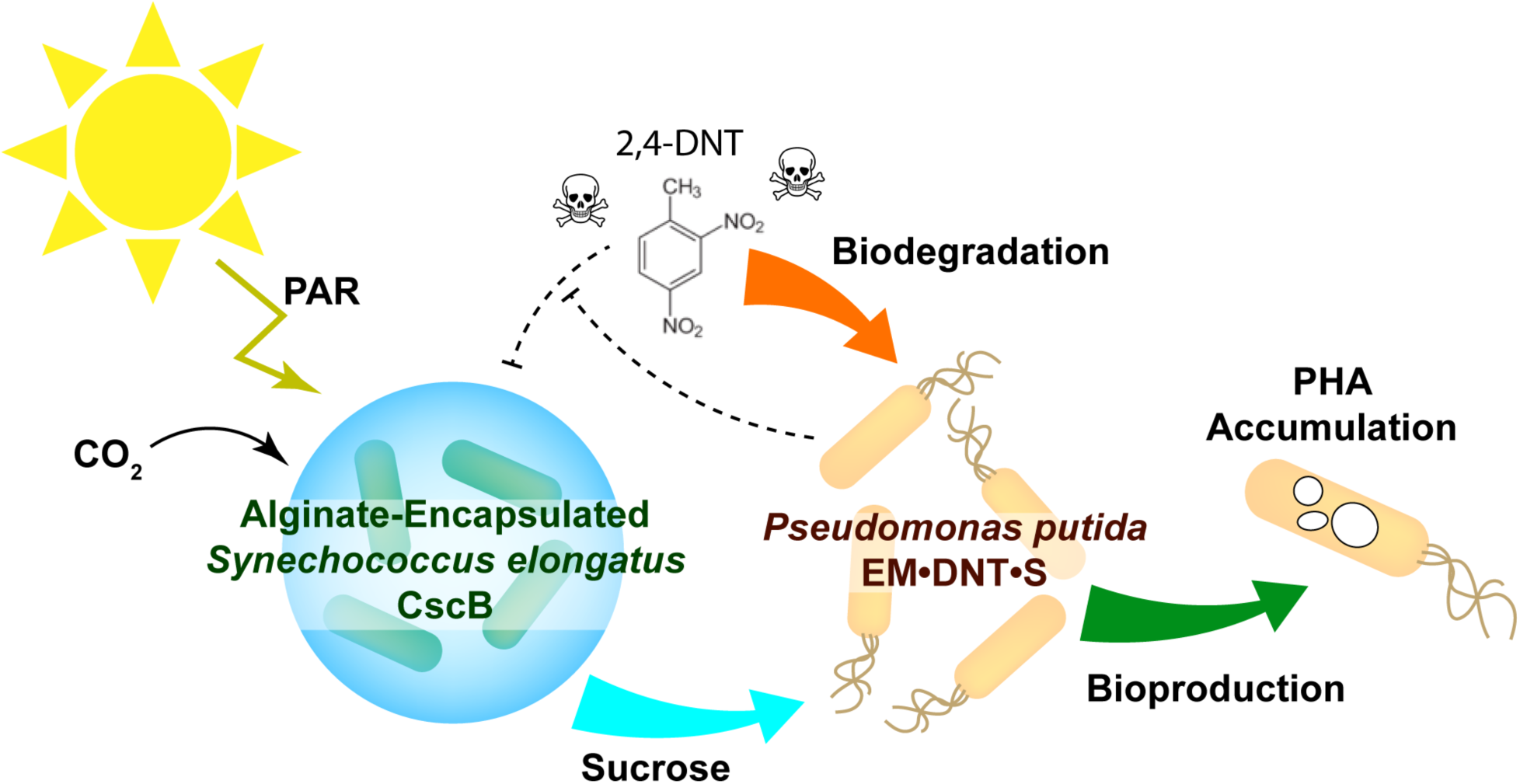
Conceptual design of the photosynthetic co-culture designed for simultaneous biodegradation and bioproduction. Alginate-encapsulated *S. elongatus* CscB is co-cultured with *P. putida* EM•DNT•S, permitting a photosynthetically-driven biodegradation of the toxin 2,4-dinitrotoluene (2,4-DNT) while simultaneously producing the bioplastic polyhydroxyalkanoate (PHA). *S. elongatus* CscB embedded within alginate beads fix carbon dioxide via the Calvin-Benson Cycle using photosynthetically active radiation (PAR). Fixed carbon is converted into sucrose that is exported through heterologous sucrose permease (CscB) in to the culture medium. Exported sucrose can be consumed by *P. putida* EM•DNT•S for growth and other metabolism, including the degradation of the toxin 2,4-DNT. Additionally, the *P. putida* EM•DNT•S can convert and store sucrose as PHA, making this system multifunctional.

## Results

### Alginate-encapsulated *S. elongatus* CscB can tolerate 2,4-DNT at higher concentrations than planktonic cultures

As 2,4-DNT is known to be highly toxic to a range of biological organisms (42, 43), we first characterized the effect of this industrial byproduct on the growth and viability of cultures of *S. elongatus* CscB. We measured culture density and the level of chlorophyll a (Chl a) in planktonic *S. elongatus* CscB cultures in the presence of increasing concentrations of 2,4-DNT (Figure 2AB). Even the lowest concentration of 2,4-DNT examined (8 µM) caused a ∼60% growth impairment in planktonic cyanobacterial cultures (Figure 2A), and a near cessation of growth in the first 48 h was observed at 2,4-DNT concentrations ≥ 15 µM (Figure 2A). The toxicity of this compound at these concentrations is highly relevant as leachates from soil contaminated with TNT and its breakdown products have been recorded as high as 98 µM (44). Chl a concentration is commonly measured as a proxy for physiological stress in cyanobacteria as chlorophyll a concentrations are strongly influenced by a variety of stress conditions (45–47). Cultures at concentrations of 2,4-DNT ranging from 31 μM to 125 μM exhibited an overall loss of Chl a (Figure 2B). This progressive loss of Chl a aligned with visual chlorosis and bleaching of these cultures; this observation was consistent at multiple 2,4-DNT concentrations and at higher starting culture densities (Figure S1 in the Supplemental Materials). These preliminary experiments indicated that planktonic cyanobacteria are highly sensitive to even low concentrations of 2,4-DNT, which would complicate their ability to be engineered to directly degrade this nitroaromatic. Further, the viability loss of planktonic *S. elongatus* CscB would need to be mitigated for any co-culture applications targeting the remediation of these compounds.

**Figure 2:**
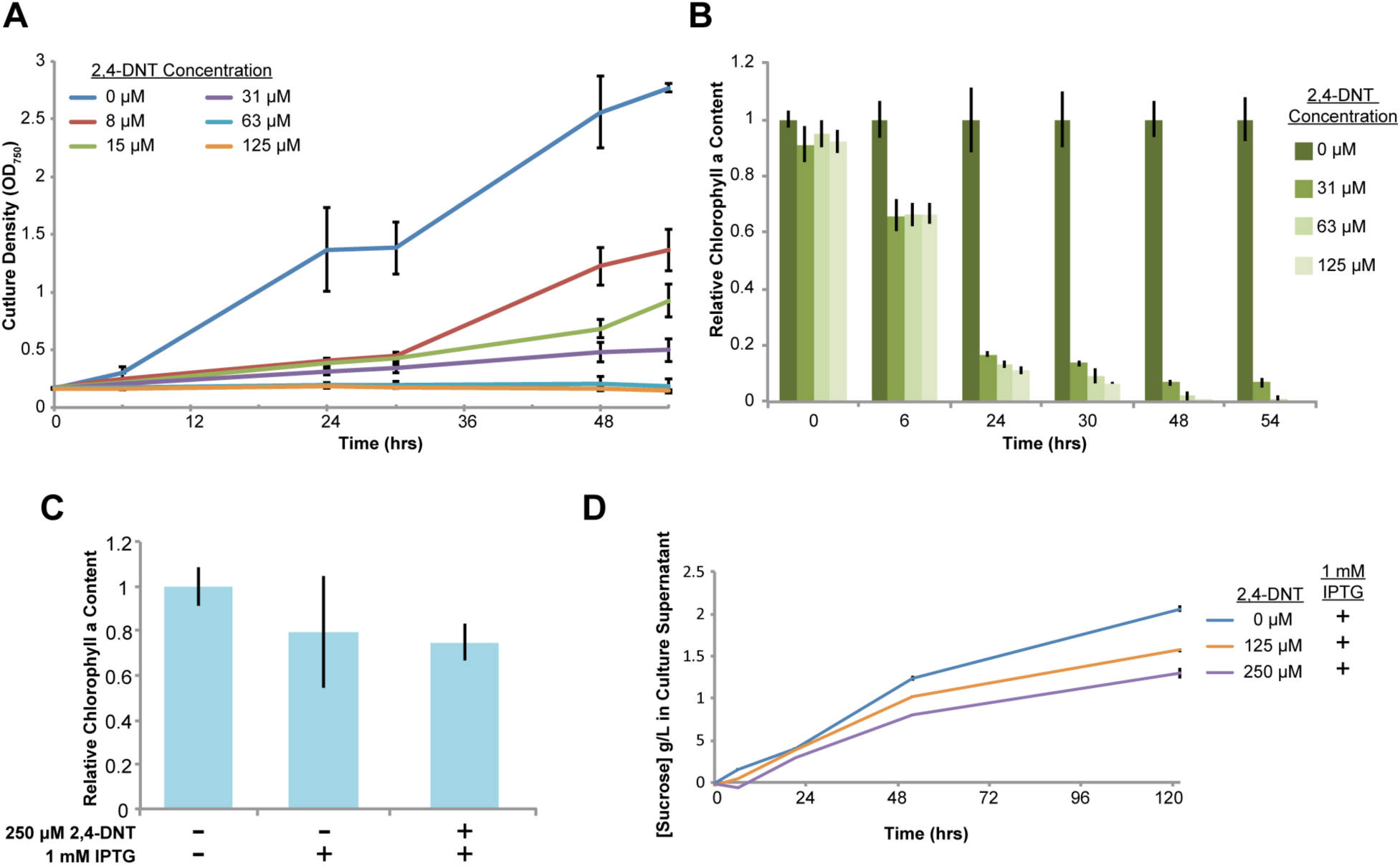
Toxic effects of 2,4-DNT exposure on planktonic and alginate-encapsulated *S. elongatus* CscB. (A) Varied concentrations of 2,4-DNT were added to planktonic cultures of *S. elongatus* CscB at OD_750_ = 0.2 and cell growth was monitored via OD_750_ for 54 h. (B) Chlorophyll a content of cultures in (a) were measured and normalized to control cultures with no added 2,4-DNT. (C) Chlorophyll a content of alginate-encapsulated *S. elongatus* CscB cells was measured after incubation for 5 days in M3 media. IPTG and/or 250 µM 2,4-DNT were added as indicated and chlorophyll a values were normalized to control cultures without these additives. (D) Total sucrose concentration in the culture supernatant of induced (1 mM IPTG) alginate-encapsulated *S. elongatus* CscB while grown with 2,4-DNT at either 0 μm, 125 μM, or 250 μM. For the experiments shown in (A-C), the mean values for *n* = 3 are indicated, and error bars represent standard deviations; for the experiment indicated in (D), the mean values for *n* = 3 with 3 technical replicates per condition are indicated, and error bars represent standard deviations.

In previous work (21), we utilized alginate hydrogel encapsulation of *S. elongatus* CscB to stabilize a co-culture with *Halomonas boliviensis* under prolonged nitrogen stress conditions. Encapsulation has been used for the immobilization of a variety of cell types, and has often led to increased stress tolerance, cell longevity, and metabolic flux toward target end products (48–53). Alginate-encapsulated *S. elongatus* CscB cells did not exhibit chlorosis in the presence of 2,4-DNT, even when the concentration was raised to 250 μM, near the solubility limit for this compound (Figure S2 in the Supplemental Materials). We then exposed encapsulated *S. elongatus* CscB cells to 2,4-DNT at 250 μM for 7 days while simultaneously inducing expression of the CscB exporter. Chl a was extracted from the beads and measured via spectrophotometry (Figure 2C). The data show that while the induction of *cscB* expression leads to a slight decrease in relative Chl a levels, the addition of 2,4-DNT to alginate-encapsulated cyanobacteria did not further decrease Chl a concentration. Similarly, the Chl a concentration per cell was maintained at a level similar to that of planktonic cells grown under our laboratory conditions (Figure S3 in the Supplemental Materials). Finally, we measured sucrose export rates from IPTG-induced, encapsulated *S. elongatus* CscB in the presence of increasing 2,4-DNT concentrations (Figure 2D). Sucrose export was maintained for multiple days despite exposure to 125 or 250 µM 2,4-DNT. Altogether, alginate encapsulation appears to stabilize *S. elongatus* when exposed to high levels of 2,4-DNT over long time periods.

### Engineering *P. putida* EM173 for sucrose consumption and evaluation of growth parameters in the presence of alginate-encapsulated *S. elongatus* CscB

We next set to construct strains of *P. putida* strains (i) that can utilize sucrose as the only carbon source and (ii) capable of degrading 2,4-DNT. *P. putida* does not normally utilize sucrose as a carbon substrate. The specific strains we used are derivatives of the genetically-tractable, prophage-less *P. putida* strain EM173 (54). To enable sucrose consumption by *P. putida* EM173, we first transformed this strain with plasmid pSEVA221-*cscRABY* (55, 56), bearing the sucrose utilization genes from *P*. *protegens* Pf-5 (Figure 3A). Specifically, this plasmid constitutively expresses genes encoding a sucrose hydrolase (CscA, PFL_3237) and a sucrose permease (CscB, PFL_3238), along with a cognate transcriptional regulator (CscR, PFL_3236). The introduction of pSEVA221-*cscRABY* into strain EM173 (giving rise to *P. putida* EM·S) enabled catabolism of sucrose and bacterial growth from the disaccharide. Separately, a synthetic mini-Tn*7* transposon, carrying the functions required for 2,4-DNT degradation in *Burkholderia* sp. R34 (Figure 3A) was constructed as described elsewhere (37). The *dnt* gene cluster in this transposon was delivered into the unique att·Tn*7* site within the chromosome of *P. putida* EM·S, resulting in a stable, engineered strain designed for 2,4-DNT degradation and sucrose consumption (*P. putida* EM·DNT·S). To test for successful catabolism of sucrose by these strains, we performed an initial characterization of sucrose catabolism in an experiment in which M9 minimal medium with 20 g/L sucrose was inoculated with either *P. putida* EM·S or *P. putida* EM·DNT·S at an optical density measured at 600 nm (OD_600_) = 0.1. These cultures were followed over the course of 24 h, monitoring both culture density (OD_600_) and soluble sucrose concentrations (Figure 3B). Both *P. putida* EM·S or *P. putida* EM·DNT·S grew exponentially over the course of the first 9 h post-inoculation and by 24 h had reached final densities of OD_600_ = 4.3 and 6 respectively. During this time, sucrose concentration had declined in a near linear fashion from 20 g/L to < 3 g/L at 24 h. As sucrose was the only added carbon source and we could clearly observe the consumption of sucrose over time, these data definitively demonstrate that the pSEVA221-*cscRABY* vector enabled sucrose utilization in these two strains.

We cultured our doubly-modified *P. putida* EM·DNT·S in the presence of a range of sucrose concentrations to determine if this strain is capable of growing on a minimal medium designed for cyanobacterial growth with sucrose as a sole carbon source. For this purpose, we generated a phosphate buffered minimal medium derived from BG-11, herein referred to as M3 medium (Table S1 in the Supplemental Material). To gauge the growth capacity of *P. putida* EM·DNT·S under these conditions, we inoculated M3 medium that did not contain a carbon source, and incubated cells overnight. This promoted acclimation to the medium and depletion of internal carbon storage compounds that could confound the accurate determination of growth in the M3 medium. These cells were washed with fresh M3 medium before being inoculated into culture flasks with a range of sucrose concentrations (0 g/L to 10 g/L) and growth was tracked for 54 h (Figure 3C). Bacterial growth was evident at sucrose concentrations ranging from 1.25-10 g/L (Figure 3C), though carbon may have been growth-limiting at concentrations lower than 2.5 g/L (Figure 3C). These results confirm that the heterologous expression of the *cscRABY* genes from *P*. *protegens* Pf-5 is sufficient to confer sucrose utilization on *P. putida* in our background strain bearing the 2,4-DNT degradation gene cluster, and indicated that the engineered *P. putida* strain can grow in the cyanobacterial M3 medium? setting the basis for conducting co-cultures.

### Growth of engineered *P. putida* strains is supported by sucrose-rich cyanobacterial exudates in a synthetic consortium system

We next explored how the engineered *P. putida* strains (*P. putida* EM·S and *P. putida* EM·DNT·S) behaved in co-culture. The *P. putida* strains, grown overnight in M3 medium with 20 g/L sucrose, were inoculated at OD_600_ ∼ 0.1 into culture flasks containing either empty alginate beads or alginate beads with encapsulated *S. elongatus* CscB (Figure 3D). In the first 48 h, both *P. putida* strains the OD_600_ continued to increase in all flasks, despite the absence of a carbon source in the flask containing empty alginate beads. Internal stores of carbon likely drove this residual growth in *P. putida*. However, these carbon stores appeared to be depleted after this period of time as the optical density of the cultures containing empty alginate beads declined over the next 24 h (Figure 3D). At 96 h, the culture supernatant was removed from all co-cultures, leaving the *S. elongatus* CscB alginate beads in place. Fresh medium was added and the co-culture densities were tracked for another 120 h (Figure 3D). Only the *P. putida* strains in flasks with alginate beads containing *S. elongatus* CscB showed signs of regrowth after the medium exchange, with OD_600_ increasing from 0.1 at 96 h to 0.5-0.6 at 216 h. These results demonstrate the *P. putida* strains tolerate and can utilize the exudates from the beaded *S. elongatus* CscBs, and are not significantly utilizing the alginate beads as a carbon source. These results are in agreement with previous work that utilized *P. putida* EM·S and *S. elongatus* CscB (17), indicating that these species are compatible under light-driven co-culture conditions.

### Biotransformation of 2,4-DNT by engineered *P. putida* EM*·*DNT*·*S in both monoculture and co-culture

We next examined the functionality of the 2,4-DNT degrading gene cluster in *P. putida* monocultures in M3 medium supplemented with 2 g/L sucrose as the sole carbon source. The *dnt* cluster is of significant interest to numerous research groups due to the fact that it is an actively evolving pathway (57, 58) and its unique biological processing of 2,4-DNT (28). This enzymatic pathway enables the oxidative degradation of 2,4-DNT, relying on two key successive dioxygenations of the 2,4-DNT substrate mediated by DntA and DntB (Figure 4A). 2,4-DNT is first dioxygenated by DntA to form 4-methyl-5-nitrocatechol (4M5NC), a compound with a strong absorption peak at 420 nm (57). 4M5NC is the substrate of DntB, which catalyzes another oxygenation reaction, transforming 4M5NC into 2-hydroxy-5-methylquinone (2H5MQ), an intermediate with an absorption peak at 485 nm (57). 2H5MQ is then processed by a number of additional genes encoded in the pathway allowing for the complete mineralization of 2,4-DNT (Figure 4A) (34, 36).

**Figure 4:**
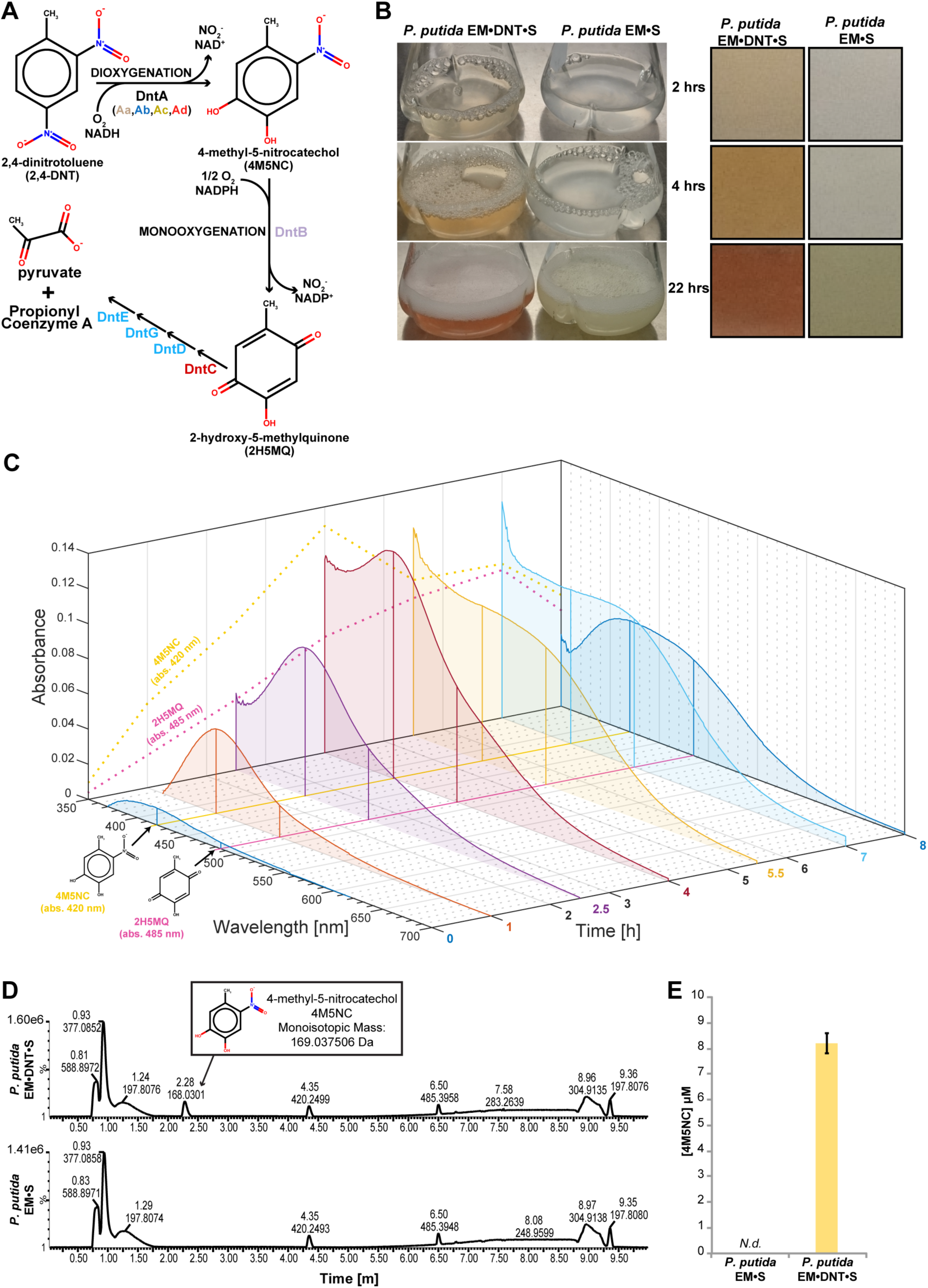
2,4-DNT biotransformation in monocultures of engineered *P. putida*. (A) Exogenous pathway for the oxidative degradation of 2,4-DNT. (B) (Left) Representative culture images of *P. putida* EM•DNT•S and *P. putida* EM•S grown over 22 h in M3 medium with 20 g/L sucrose in the presence of 250 μM 2,4-DNT. (Right) Zoom-in of cultures demonstrating characteristic changes in pigmentation of culture supernatants. (C) Averaged spectral signatures (*n*=3) of supernatants from *P. putida* EM•DNT•S cultures grown as in M3 medium with 2 g/L sucrose in the presence of 250 μM 2,4-DNT. Samples were monitored via scanning spectrophotometry at the indicated time points. (D) Representative LC/MS elution profile of *P. putida* EM•DNT•S (top) and *P. putida* EM•S (bottom) supernatant after 4 hours of incubation M3 medium with 2 g/L sucrose and 250 μM 2,4-DNT. 4M5NC elutes from the column after 2.3 min in negative ion mode. (E) Quantification of 4M5NC in supernatants of both *P. putida* EM•S and *P. putida* EM•DNT•S monocultures at the 4 h via LC/MS. *P. putida* EM•S strain did not generate any detectable amount of 4M5NC, while the *P. putida* EM•DNT•S strain accumulated 4M5NC.

We examined the transformation of 2,4-DNT by either *P. putida* EM·S or *P. putida* EM·DNT·S strains over time (Figure 4B-E). Flasks containing *P. putida* EM·S or *P. putida* EM·DNT·S were inoculated at OD_600_ ∼ 0.1 and 2,4-DNT was added at 250 μM to the cultures. The supernatant of the *P. putida* EM·DNT·S cultures was visibly changed during the experiment (Figure 4B), and these spectroscopic shifts were consistent with the accumulation of intermediates of 2,4-DNT breakdown through the exogenous oxidative pathway (Figure 4A). As soon as 2 h after addition of 2,4-DNT to the culture medium at 250 μM, the supernatant turned yellow (Figure 4B), which is consistent with the accumulation of the first pathway intermediate, 4M5NC (57). Four hours later, the supernatant became visibly orange (Figure 4B), suggestive of accumulation of the second intermediate, 2H5MQ (57). These changes in supernatant coloration were not observed in the control reactions with the non-degrading *P. putida* EM·S strain (Figure 4B).

We further verified that the colorimetric changes in the supernatant of *P. putida* EM·DNT·S exposed to 2,4-DNT could be attributed to breakdown of the compound through the heterologous *dnt* pathway. An 8h time-course experiment was performed with additional time points in which both strains of *P. putida* were grown in M3 medium with 2 g/L sucrose during which the cultures were sampled and the supernatant extracted. The absorbance spectra of *P. putida* EM·DNT·S supernatants exhibited a characteristic rise in an absorption peak at 420 nm over time until 4 h, at which time a second absorption peak at 485 nm became evident (Figure 4C). No defined peaks were evident in the visible wavelength absorption spectra of the non-degrading control strain (*P. putida* EM·S) (Figure S4 in the Supplemental Materials). Although spectroscopic analysis is well-supported in the literature as a metric for measuring activity of this oxidative pathway (57), a more direct method of quantification was desirable to confirm the appearance of the pathway intermediates. Hence, supernatant samples from both *P. putida* EM·S and EM·DNT·S cultures at the 4h time point were evaluated for the presence of 4M5NC by liquid chromatography coupled to mass spectrometry (LC/MS). Figure 3D shows a representative LC/MS profile from one of the *P. putida* EM·DNT·S cultures, the peak representing 4M5NC is indicated. This peak matches that of the 4M5NC standard included for quantification (Figure S5 in the Supplemental materials), and comparison to cultures of the non-degrading strain indicate that there was no 4M5NC generated by the control strain (Figure 4D-E).

To confirm specific degradation of the 2,4-DNT, culture supernatants were periodically sampled over two days. 2,4-DNT was observed to be rapidly lost from *P. putida* EM·DNT·S cultures over the course of the first 4 h, as measured by tandem gas chromatography coupled to mass spectrometry (GC-MS), and had dropped to undetectable levels by 22 h, a kinetic pattern of 2,4-DNT transformation similar to what has been reported for *Burkholderia* sp. R34 (38) and other engineered *P. putida* strains (37) (Figure S6A in the Supplemental Materials). 2,4-DNT concentrations in cultures containing the *P. putida* EM·S strain appeared to decrease at a similar rate (Figure S6A in the Supplemental Materials). This loss of 2,4-DNT may be attributed to the adsorption of the 2,4-DNT to cell surfaces, a known property of nitroaromatic compounds that makes them difficult to extract from biological substrate (33), and/or the reduction of the compound by non-specific nitroreductases. Previous work by Akkaya et al. (37) with the *P. putida* EM strain indicated that the reductive pathway (Figure S6C in the Supplemental Materials) is not likely a major contributor to 2,4-DNT transformation. We assessed our supernatant samples for the presence of 2-amino-4-nitrotoluene/4-amino-2-nitrotoluene (2A4NT/4A2NT), intermediates in the non-specific reductive pathway (Figure S6D in the Supplemental Materials). These compounds were detected in minor amounts (< 3 μM) in both the *P. putida* EM·DNT·S and *P. putida* EM·S cultures (Figure S6D in the Supplemental Materials). While it is difficult to quantify the flux attributable to the reductive pathway, the accumulation of 2A4NT/4A2NT compounds was reduced by 22% in *P. putida* EM·DNT·S cultures compared to the *P. putida* EM·S (Figure S6D in the Supplemental Materials).

### Degradation of 2,4-DNT by a synthetic consortium of *P. putida* EM*·*DNT*·*S and alginate-encapsulated *S. elongatus* CscB and long-term culture potential

We introduced 2,4-DNT into co-cultures of *P. putida* and encapsulated *S. elongatus* CscB. Both the *P. putida* EM·S and *P. putida* EM·DNT·S strains were able to grow with the beaded *S. elongatus* CscB in the presence of 250 μM 2,4-DNT (Figure 5A). Cultures containing only encapsulated *S. elongatus* CscB or empty alginate beads served as controls for both optical density measurements as well as any potential 2,4-DNT adsorption. GC-MS analysis for 2,4-DNT content of these cultures demonstrated that cultures containing *P. putida* or *S. elongatus* cells removed 2,4-DNT from the culture (Figure 5B), in line with previous results. In contrast, 2,4-DNT concentrations did not decrease in the flasks containing only empty beads (Figure 5B). Co-cultures containing *P. putida* EM·DNT·S demonstrated a similar color change in the culture supernatant to that of *P. putida* EM·DNT·S monocultures, indicating that the exogenous pathway retained its function in the consortium system. These results demonstrate that this system can provide a directed method for the photosynthetically-driven degradation of 2,4-DNT. Subsequent LC-MS testing of co-culture supernatants shortly (1 h) and 24 h after inoculation of the co-culture with 2,4-DNT revealed that, as in previous monoculture experiments, the *P. putida* EM·DNT·S co-cultures accumulated 4M5NC (Figure 5C). This observation, again, indicates that 2,4-DNT is being degraded via the oxidative pathways in these co-cultures.

**Figure 5:**
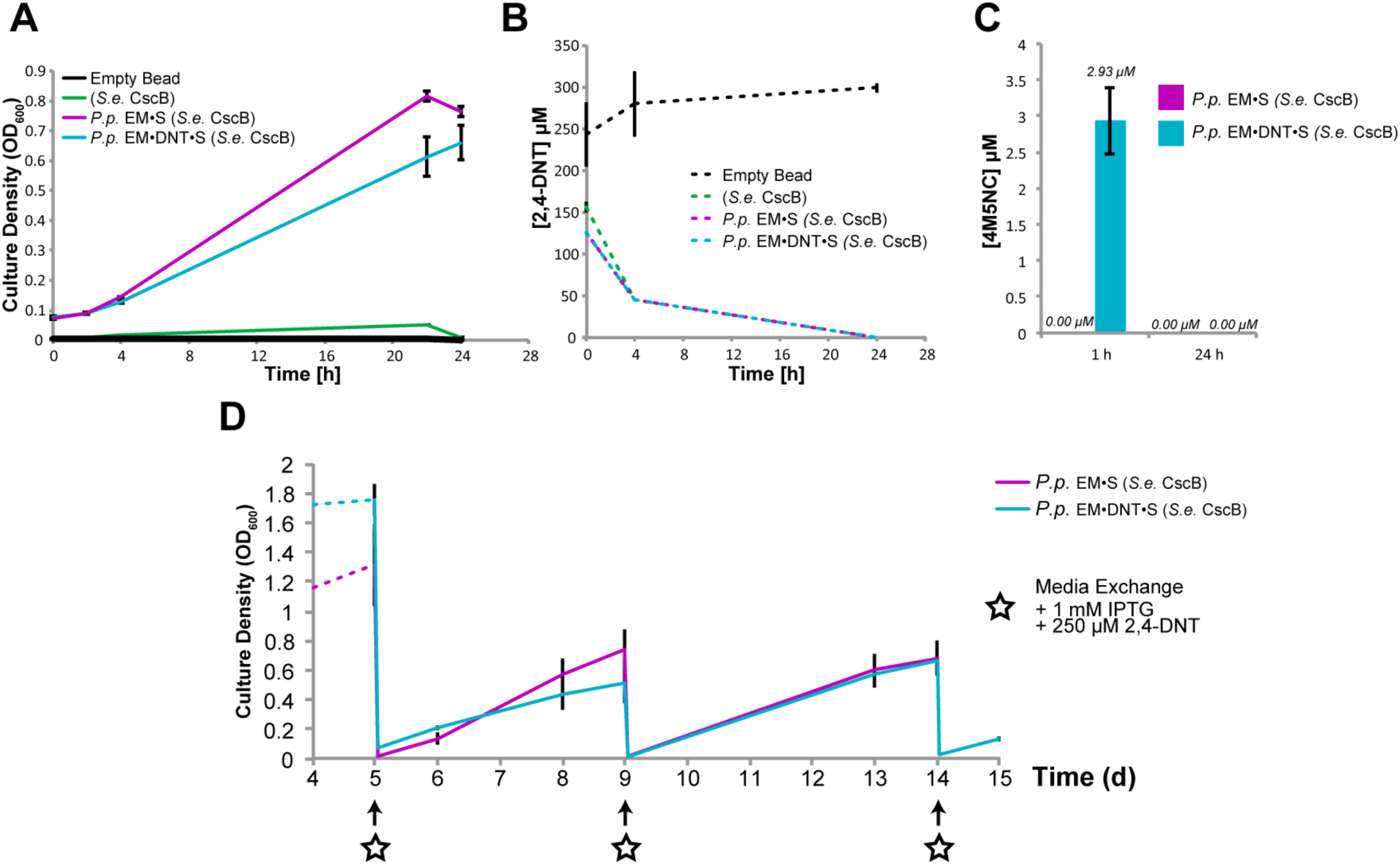
Biotransformation of 2,4-DNT in *S. elongatus/P. putida* co-culture. (A) Growth of indicated planktonic *P. putida* strains in co-culture with alginate-encapsulated *S. elongatus* CscB with 250 μM 2,4-DNT. (B) GC/MS analysis of supernatants from 24 hr co-cultures demonstrates transformation of 2,4-DNT. 2,4-DNT concentrations dropped below detectable levels by 24 hours post inoculation. (C) LC/MS quantification of 4M5NC content in co-cultures 1 h or 24 h after 2,4-DNT addition. *P. putida* EM•DNT•S cultures accumulated 4M5NC that is been fully degraded by 24 h. (D) Optical density of a long-term co-culture of *P. putida* strains with encapsulated *S. elongatus* CscB. Media is exchanged a at each indicated timepoint (star), where the culture supernatant containing planktonic *P. putida* is removed and fresh media is added back to the culture along with 1 mM of IPTG and 250 μM 2,4-DNT. Residual *P. putida* cells grow following each media exchange. For (A-C), *n* = 3, error bars represent standard deviations. For (D), *n* = 14, error bars represent standard deviations.

We initiated a longer duration co-culture in which we inoculated both strains of *P. putida* into culture flasks with encapsulated *S. elongatus* CscB cells and followed the cultures for 15 days, exchanging the culture supernatant every 4-5 days with fresh medium containing 250 μM 2,4-DNT (Figure 5D). Regrowth of *P. putida* strains following backdilution indicate stable repopulation, allowing for continual degradation of the 2,4-DNT (Figure 5D). The total mass of 2,4-DNT cleared by the *P. putida* EM·DNT·S strain over this period amounts to ca. 4 mg.

### Simultaneous 2,4-DNT biodegradation and PHA bioproduction by engineered strains in a synthetic consortium

*P. putida* accumulates PHA as a carbon storage, particularly under nitrogen-depleted conditions (17), providing the opportunity to both bioremediate 2,4-DNT and simultaneously produce a valuable byproduct. *P. putida* cells were inoculated into the encapsulated and induced *S. elongatus* CscB culture flasks at OD_600_ ∼ 0.5 with 250 μM 2,4-DNT (Figure 6A). These flasks contained either M3 or M3-N medium, the latter of which has a reduced nitrogen content (2mM NH_4_Cl) (Table 3.S1). Twenty four hours after cycling into the nitrogen deplete medium, cell culture was harvested, dried, and processed for quantification of methyl esters of alkanoic acids by high-pressure liquid chromatography (HPLC; see Methods). Processing of these samples showed that both *P. putida* EM·S and *P. putida* EM·DNT·S co-cultures in nitrogen deplete medium accumulated 4.9 and 5.1 mg PHA/L of culture, resulting in polymer contents of 22 and 23.4 mg of PHA/g cell dry weight, respectively.

**Figure 6:**
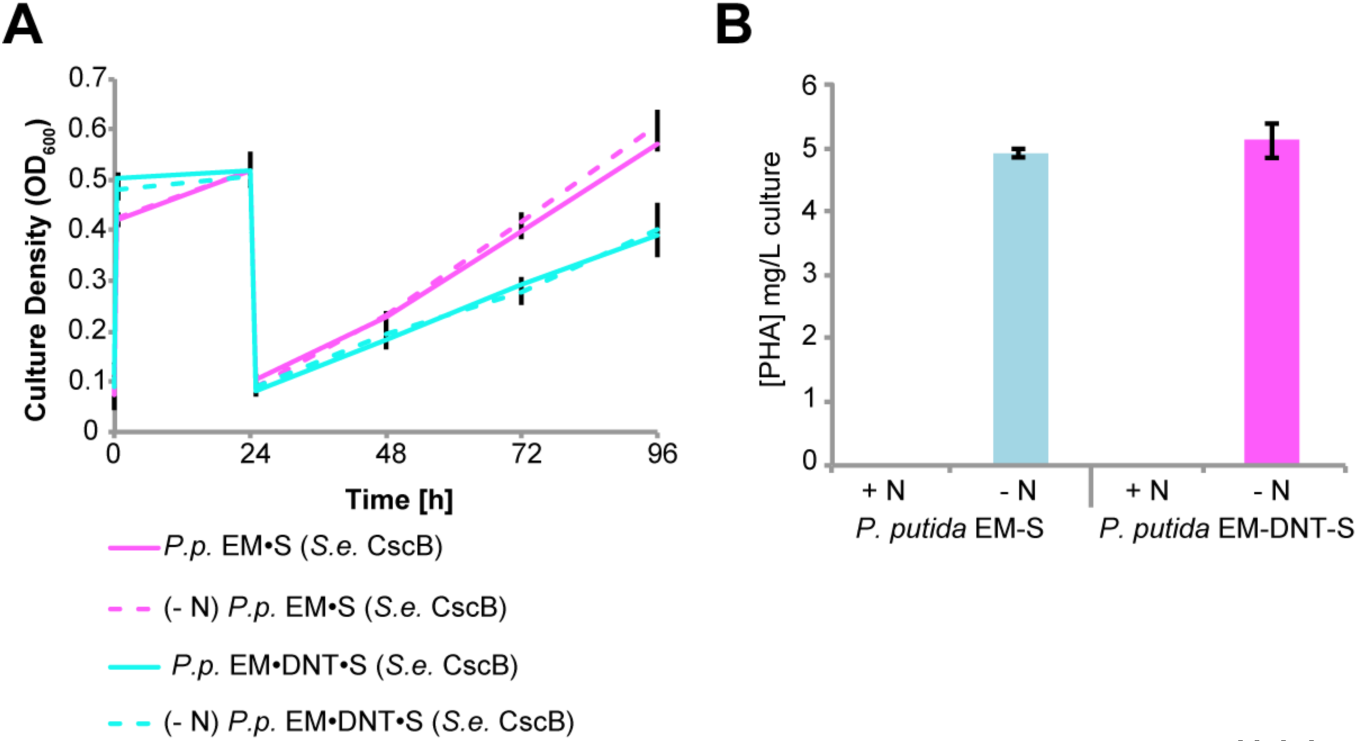
PHA accumulation in *P. putida* bioremediating co-cultures. (A) Co-cultures of beaded *S. elongatus* CscB and *P. putida* strains in the presence of 2,4-DNT with the standard (17.6 mM) or reduced concentration (2 mM) of nitrate. Nitrogen scarcity triggers *P. putida* to allocate excess carbon into polyhydroxyalkanoate (PHA). (B) Quantification of PHA extracted from the *P. putida* biomass of co-cultures 24 h after inoculation into nitrogen replete (+ N) or nitrogen deplete (-N) medium. For (A) and (B), *n* = 3; error bars represent standard deviations.

## Discussion

In this work, we demonstrate metabolic division of labor within an artificial microbial co-culture consisting of engineered strains of *S. elongatus* PCC 7942 and *P. putida* EM173. In this consortium, *S. elongatus* CscB produces soluble sugars using light and CO_2_ as inputs, providing sufficient organic compounds to promote the growth of co-cultivated strains of *P. putida.* In turn, we demonstrate that *P. putida* simultaneously degrades the environmental pollutant 2,4-DNT while producing the bioplastic precursor, PHA.

We have shown that encapsulation of *S. elongatus* CscB cells in alginate hydrogel beads allowed the cyanobacteria to persist in the presence of 2,4-DNT (Figs. 3.S1-S2 in the Supplemental Materials) and maintain consistent production of sucrose after culture back dilution. Encapsulated cyanobacteria tolerated concentrations of 2,4-DNT 5 times higher than could be tolerated by planktonic cyanobacterial cultures and did so without significantly changing chlorophyll a content (Figure 2BC). While there is precedence for the improved resilience of cells that are encapsulated within hydrogels (21, 59), the mechanism by which this occurs has yet to be elucidated. Future work to investigate how encapsulation modulates cyanobacterium stress response to nitroaromatic compounds could yield genetic targets that could be modified to bolster the resilience of the cyanobacteria without the need for mechanical encapsulation.

*P. putida* is a gram-negative soil bacterium that has recently gained significant attention as a *chassis* for industrially-relevant synthetic biology. This is in part thanks to the full sequencing (40) and subsequent generation of *P. putida* strains (*e.g., P. putida* EM173 used in this work) with reduced genomes that demonstrate enhanced expression of heterologous proteins (54). *P. putida*’s metabolic diversity and high tolerance of oxidative stress make it an ideal model organism for studying toxin remediation as well as bioproduction of added-value compounds (41). The introduced oxidative 2,4-DNT degradation pathway (Figure 4A) is chromosomally integrated in this strain. While this pathway avoids generating highly reactive intermediates with hydroxylamino groups, the proteins in this pathway are not yet fully optimized for this new substrate (38, 57), leading to the production of oxidative stress in the *Burkholderia sp.* R34 from which the pathway originates (38). This oxidative damage is thought to contribute to an increased rate of mutation and fosters a more rapid evolution of this strain to combat oxidative stress (38). While *P. putida* has also been shown to experience increased oxidative stress when actively utilizing this pathway, this species does not exhibit the same rate of DNA damage and mutation (37). Thus, the physiological properties of *P. putida* allow for more efficient use of an imperfect 2,4-DNT degradation pathway. Evolving or engineering this pathway toward increased specificity for 2,4-DNT could allow for improved kinetics and reduced ROS generation. Conversely, the lower substrate specificity might allow this pathway to be redirected toward the degradation of other types of nitroaromatic pollutants from other industrial processes (23).

2,4-DNT as a compound represents a significant contaminant that poses a bioremediation challenge. 2,4-DNT in its solid state is very stable in the environment as it is not bioavailable, where it often remains due to its low solubility. Once solubilized, 2,4-DNT can then be reduced to highly reactive hydroxylamino intermediates that damage cellular machinery and DNA as chemical adducts. Solid state 2,4-DNT has a high stability and low solubility, allowing this compound to persist in the environment and slowly disperse into surface and ground water. Furthermore, reduced derivatives of this compound do not readily mineralize in the environment and persist for extended periods of time in the soil (33). The longevity of this compound and its derivatives requires a long-term sustainable solution to remediate previous areas of contamination, as well as reduce contamination from industrial practices that are actively producing 2,4-DNT as a by-product. Previous works that propose alternative biologically based methods of degrading 2,4-DNT are limited either by the necessity to supply bioavailable carbon (60) or produce new biological substrate to perform the degradation (61).

Co-cultures of the 2,4-DNT degrading *P. putida* strain (*P. putida* EM·DNT·S) were successfully grown while solely supported by the fixed carbon provided by the encapsulated *S. elongatus* CscB. Furthermore, these cultures were able to successfully cycle over the two weeks as a demonstration of this system’s stability and potential for long-term sustainable degradation of 2,4-DNT. While we demonstrate intermittent cycling of media in flask-based cultures here (Figure 5D), it is possible to conceive of a photobioreactor that would allow for continual introduction of 2,4-DNT-contaminated wastewater. Here, the full media exchanges associated with long-term flask-based co-cultures allowed us to demonstrate the ability of these co-cultures not only to remediate this compound but also produce the bioplastic precursor polyhydroxyalkanoate (PHA).

While sustainable production of PHA has been pursued in other contexts, this is, to our knowledge, the first report in which PHA formation has been concurrent with the degradation of a xenobiotic compound. In comparison to an independent report that aimed to optimize PHA production from batch cultures of *S. elongatus-P. putida* (17), we achieved a lower specific productivity in cultures simultaneously degrading 2,4-DNT. Increased productivities of PHA might be achieved by modifying total nitrogen supplied, duration of nitrogen starvation, or concentration of 2,4-DNT. Of note, we observed no appreciable difference in *P. putida* growth rates between co-cultures in nitrogen replete or in low nitrogen (2mM nitrate). This raises the possibility that *P. putida* may be able to utilize an unknown cyanobacterial by-product as a nitrogen source, and additional optimization may be required to fully activate PHA production pathways.

One question that arose as part of this work was whether the presence of *P. putida* EM·DNT·S provides a protective effect to *S. elongatus* CscB in co-cultures fed with 2,4-DNT contaminated medium. If this were the case, it would shift this relationship from a commensal to a more mutualistic relationship where each species benefits from the presence of the other. Pursuing longer-term cultures with even more rigorous exposure to 2,4-DNT could reveal whether *P. putida* EM·DNT·S might provide such a protective effect.

We show that these artificial co-cultures are not only capable of utilizing media contaminated with a toxic xenobiotic, but also of producing the bioplastic PHA. A scaled version of this system could hypothetically take wastewater effluent from industrial sources contaminated with 2,4-DNT, remediate the water allowing it to be utilized for other functions, and provide a mechanism by which PHA could be produced. The more immediately relevant take-away from this work is that photosynthetic co-cultures utilizing *S. elongatus* CscB are flexible both in its partnerships with other microbes as well as the intended functionality of the system. This creates opportunities for more advanced and complex co-cultures with new functions and constituents that we hope will not only provide solutions to modern conflicts, but also inform us on how nature of symbiotic relationships develop in the natural world.

## Methods

### Bacterial strains and culture conditions

Planktonic *S. elongatus* strains were grown as previously described (21) within a Multitron Pro (Infors) photobiological shaker at 32°C, 2% CO2, and a constant illumination of ∼70 μmol m^-2^ s^-1^ (15W Grow-Lux; Sylvania). Selection for the genomically-integrated *cscB* cassette was maintained during with the addition of 12.5 ug/mL chloramphenicol to the medium. *P. putida* strains were constructed as reported previously (55, 57). During routine cultivation, *P. putida* strains with an integrated *dnt* degradation cassette were maintained with 25 μg/mL gentamicin while strains transformed with plasmid pSEVA221-cscRABY were maintained with 50 ug/mL kanamycin. LB cultures were at 32°C in a Multitron (Infors) incubator with rotary agitation at 150 rpm. For co-culture experiments, *P. putida* cultures grown in LB were resuspended in M3 medium (see Table S1 in the Supplemental Material) with either 20 g/L or 2 g/L sucrose, as indicated. These cultures were then used as inoculum for subsequent experiments. Antibiotic selection was omitted for all co-culture experiments.

### Encapsulation of *S. elongatus* CscB in alginate beads

Alginate encapsulation was performed as previously described (21) with minor adjustments. Briefly, planktonic *S. elongatus* CscB cells grown in BG-11 medium + 1g/L HEPES (pH 8.3; Sigma) were harvested at an optical density at 750 nm (OD_750_) of 2.0 via centrifugation at 3,500×g for 30 min and concentrated in 3 mL of sulfur-free BG-11. Concentrated cells were mixed into a sterile and degassed volume of 3% (wt/vol) sodium alginate at a final OD_750_ = 5.0; a ∼2.75% (wt/vol) sodium alginate-*S. elongatus* CscB suspension. In a sterile biosafety cabinet, a vertically-oriented syringe pump (KD Scientific, Holliston, MA) was used to dispense the solution dropwise through a 30 G needle into a ≥20-fold larger volume of 20 mM BaCl_2_. The drops traveled ∼35 cm from needle to the slowly stirred BaCl_2_ solution and were allowed to cure in the solution for at least 20 min. Solidified beads were rinsed once with BG-11 medium and incubated overnight in M3 medium (minus the 100 mM NaCl). To acclimate cells within the alginate and precipitate residual Ba^2+^, beads were transferred through a series of media washes in following days. The first day post-encapsulation, beads were rinsed and resuspended in fresh M3 medium (without NaCl), transferred to 250-mL baffled Erlenmeyer flasks, and placed into a Multitron Pro (Infors) and shaking (125 rpm), as above for 24 h. Subsequently, M3 medium was exchanged, and the alginate beads were portioned in ∼10-mL aliquots into 125-mL baffled Erlenmeyer flasks. To acclimate cells to final salt concentrations, medium was replaced with M3 with only 25 mM NaCl for 24 hours, then M3 with 50 mM NaCl for 24h, and finally to complete M3 (with 100mM NaCl). The medium was subsequently refreshed daily for 3-5 days prior to initiation of experiments. In the cases where the beads would be utilized in co-culture, *cscB* expression was induced with the addition of +1mM IPTG 24 hours prior to addition of *P. putida* cells. Media was exchanged a final time coincident with addition of *P. putida*.

### Analytical methods

Culture optical densities were measured with a Genesys 20 (Thermo Fisher Scientific, Waltham, MA) spectrophotometer. Planktonic *S. elongatus* CscB was measured at OD_750_, and both co-culture experiments and *P. putida* monocultures were measured at OD_600_. This spectrophotometer was also used to measure chlorophyll a concentrations of planktonic *S. elongatus* cultures, as in (62). For chlorophyll a measurements in alginate-encapsulated *S. elongatus*, 1 mL of pre-chilled methanol was added to four alginate beads, with 3 technical replicates. Beads were vortexed and incubated at 4°C for 30 min in the dark to extract chlorophyll. Absorbance of the supernatant was measured in cuvettes at 720 nm and 665 nm to calculate the final chlorophyll a concentration as previously described (62).

Sucrose concentration of beaded cultures was measured as indicated previously (21). Briefly, 1 mL of culture supernatant was withdrawn at indicated time points, cells were pelleted at 17,000×g for 10 min, and supernatant sucrose was quantified via sucrose/D-glucose assay kit (Megazyme, Bray, Ireland) with 3 technical replicates for each sample (21).

Culture supernatant spectra were measured using a DU800 Spectrophotometer, (Beckman Coulter, CA, USA). Cell-free culture supernatant was obtained by centrifugation for 10 min at 17,000×g. The supernatant was then transferred to a cuvette for spectral analysis.

GC/MS and LC/MS detection and quantitation were performed at the Michigan State University Mass Spectrometry and Metabolomics Core. 2,4-DNT measurements were made using an Agilent 5975 GC/single quadrupole MS. Culture samples were centrifuged at 17,000×g for 10 min and 10 μL of cell free supernatant were removed to a new tube. 50 μL of ethyl acetate was added to the supernatant and allowed to incubate at room temperature for 30 min. The 50 μL of ethyl acetate was then transferred to a GC vial and injected (injection volume 1 μL). Samples were separated with a 5% phenyl-methyl capillary column (Agilent) and measured by Mass Selective Detector (MSD). A temperature of 275°C was set for the split/splitless injector (ratio of 10:1). Helium gas was used as the carrier gas at a flow rate of 1 mL/min.

LC/MS measurements were made using a Waters Xevo G2-XS UPLC/MS/MS (Waters). Culture samples were harvested and pelleted via centrifugation (17,000×g for 10 min). A 100-μL aliquot of the supernatant was then transferred to a clean tube, lyophilized, and resuspended in 1 mL deionized water. Supernatant samples were harvested from culture and HPLC Measurements of PHA accumulation were made with a Waters 2695 HPLC. Samples were process and analyzed as previously described (21). Briefly, cells were centrifuged at 17,000×g for 10 min, the supernatant decanted, and the pellet was lyophilized. The cell biomass was dissolved in 1 mL of concentrated sulfuric acid, heated to 90 °C for 1 h, cooled to room temperature, and then diluted 100-fold with deionized water. A 20-μL aliquot was injected onto an Aminex 300-mm HPX-87H (Bio-Rad Laboratories, Hercules, CA) column, and 0.028 N H2SO4 was used as the mobile phase at a flow of 1 mL/min. The column temperature was maintained at 60 °C and UV-absorption was monitored at 210 nm. Two standards of commercial PHB (Sigma-Aldrich) were similarly treated and used for quantification purposes.

## ACKNOWLEDGEMENTS

This work was supported by the National Science Foundation (Award Number #1437657). Additional institutional and equipment support was provided by Office of Basic Energy Sciences, Office of Science, US Department of Energy (DE–FG02–91ER20021). This study was also supported by The Novo Nordisk Foundation (Grant NNF10CC1016517) and the Danish Council for Independent Research (SWEET, DFF-Research Project 8021-00039B) to P.I.N. The authors would like to thank Dr. Dan Jones, Dr. Anthony Schilmiller, and Dr. Scott Smith from the MSU Mass Spectrometry and Metabolomics Core for their assistance in data acquisition and analysis during this project, as well as Dr. Cheryl Kerfeld for the use of the HPLC. The authors are also indebted to Hannes Löwe and Katharina Pflüger-Grau (Technische Universität München, Germany) for sharing research materials.

## Author Contributions

DTF, PIN, and DCD designed and directed the project and experiments. PC designed and engineered the *P. putida* strains utilized in this work. DTF and PS performed the experiments and recorded the results. DTF, PS, PIN, and DCD analyzed and interpreted the data. DTF prepared the manuscript and figures. DTF, PS, PC, PIN, and DCD provided commentary and edits to the manuscript and figures.

## References

1. Little AE, Robinson CJ, Peterson SB, Raffa KF, Handelsman J. 2008. Rules of Engagement: Interspecies Interactions that Regulate Microbial Communities. Annu Rev Microbiol 62:375–401.

2. Saxena S. 2015. Diversity of Industrially Relevant Microbes, p. 1–11. In Applied Microbiology. Springer India, New Delhi.

3. Dolinšek J, Goldschmidt F, Johnson DR. 2016. Synthetic microbial ecology and the dynamic interplay between microbial genotypes. FEMS Microbiol Rev 1–19.

4. Goldford JE, Lu N, Bajic D, Estrela S, Tikhonov M, Sanchez-Gorostiaga A, Segre D, Mehta P, Sanchez A. 2017. Emergent Simplicity in Microbial Community Assembly. bioRxiv 474:205831.

5. Gestel J VaN, Vlamakis H, Kolter R. 2015. Division of Labor in Biofilms?: the Ecology of Cell Differentiation. Microbiol Spectr Am Soc Microbiol 1–24.

6. Tan J, Zuniga C, Zengler K. 2015. Unraveling interactions in microbial communities - from co-cultures to microbiomes. J Microbiol 53:295–305.

7. Tecon R, Or D. 2017. Cooperation in carbon source degradation shapes spatial self-organization of microbial consortia on hydrated surfaces. Sci Rep 7:1–11.

8. Nikel PI, Silva-Rocha R, Benedetti I, de Lorenzo V. 2014. The private life of environmental bacteria: Pollutant biodegradation at the single cell level. Environ Microbiol 16:628–642.

9. Wilkinson TG, Topiwala HH, Hamer G. 1974. Interactions in a mixed bacterial population growing on methane in continuous culture. Biotechnol Bioeng 16:41–59.

10. Kumar KH, Jagadeesh KS. 2016. Microbial consortia-mediated plant defense against phytopathogens and growth benefits. South Indian J Biol Sci 2:395–403.

11. Blasche S, Kim Y, Oliveira AP, Patil KR. 2017. Model Microbial Communities for Ecosystems Biology. Curr Opin Syst Biol 1–7.

12. Ortiz-Marquez JCF, Do Nascimento M, Zehr JP, Curatti L. 2013. Genetic engineering of multispecies microbial cell factories as an alternative for bioenergy production. Trends Biotechnol 31:521–9.

13. Ducat DC, Avelar-Rivas JA, Way JC, Silver PA. 2012. Rerouting carbon flux to enhance photosynthetic productivity. Appl Environ Microbiol 78:2660–8.

14. Song K, Tan X, Liang Y, Lu X. 2016. The potential of Synechococcus elongatus UTEX 2973 for sugar feedstock production. Appl Microbiol Biotechnol.

15. Du W, Liang F, Duan Y, Tan X, Lu X. 2013. Exploring the photosynthetic production capacity of sucrose by cyanobacteria. Metab Eng 19:17–25.

16. Hays SG, Yan LLW, Silver PA, Ducat DC. 2017. Synthetic Photosynthetic Consortia Define Interactions Leading to Robustness and Photoproduction. J Biol Eng 11:1–36.

17. Löwe H, Hobmeier K, Moos M, Kremling A, Pflüger-Grau K. 2017. Photoautotrophic production of polyhydroxyalkanoates in a synthetic mixed culture of Synechococcus elongatus cscB and Pseudomonas putida cscAB. Biotechnol Biofuels 1–11.

18. Smith MJ, Francis MB. 2016. A Designed A. vinelandii-S. elongatus Coculture for Chemical Photoproduction from Air, Water, Phosphate, and Trace Metals. ACS Synth Biol 5:acssynbio.6b00107.

19. Smith MJ, Francis MB. 2017. Improving metabolite production in microbial co-cultures using a spatially constrained hydrogel. Biotechnol Bioeng 114:1195–1200.

20. Li T, Li C-T, Butler K, Hays SG, Guarnieri MT, Oyler GA, Betenbaugh MJ. 2017. Mimicking lichens: Incorporation of yeast strains together with sucrose-secreting cyanobacteria improves survival, growth, ROS removal, and lipid production in a stable mutualistic co-culture production platform. Biotechnol Biofuels 10:1–12.

21. Weiss TL, Young EJ, Ducat DC. 2017. A synthetic, light-driven consortium of cyanobacteria and heterotrophic bacteria enables stable polyhydroxybutyrate production. Metab Eng 44:236–245.

22. Barry A, Wolfe A, English C, Ruddick C, Lambert D. 2016. 2016 National Algal Biofuels Technology ReviewBioenergy Technologies Office.

23. Ju K-S, Parales R E.2010. Nitroaromatic Compounds, from Synthesis to Biodegradation. Microbiol Mol Biol Rev 74:250–272.

24. Spain JC. 1995. Biodegradation of Nitroaromatic Compounds. Annu Rev Microbiol 49:2548–2523–555.

25. Yinon J. 1990. Toxicity and Metabolism of Explosives. CRC Press, Inc., Boca Raton, Florida, USA.

26. Hooker BS, Skeen RS. 1999. Transgenic phytoremediation blasts onto the scene. Nat Biotechnol 17:1999.

27. Griest WH, Stewart AJ, Vass AA, Ho CH. 1998. Chemical and Toxicological Characterization of Slurry Reactor Biotreatment of Explosives-Contaminated Soils.pdf. Oak Ridge, Tennessee, USA.

28. Symons ZC, Bruce NC. 2006. Bacterial pathways for degradation of nitroaromatics. Nat Prod Rep 23:845.

29. French CE, Rosser SJ, Bruce NC. 2001. Biotransformations of Explosives. Biotechnol Genet Eng Rev 18:171–217.

30. Bryant C, McElroy WD. 1991. Nitroreductases, p. 291–299. In Muller, F (ed.), Chemistry and Biochemistry of Flavoenzymes, Volume II, 1st ed. CRC Press, Inc., Boca Raton, Florida, USA.

31. Park W, Peña-Llopis S, Lee Y, Demple B. 2006. Regulation of superoxide stress in Pseudomonas putida KT2440 is different from the SoxR paradigm in Escherichia coli. Biochem Biophys Res Commun 341:51–56.

32. Padda RS, Wang C, Hughes JB, Kutty R, Bennett GN. 2003. Mutagenicity of nitroaromatic degradation compounds. Environ Toxicol Chem 22:2293–2297.

33. Achtnich C, Sieglen U, Knackmuss HJ, Lenke H. 1999. Irreversible binding of biologically reduced 2,4,6-trinitrotoluene to soil. Environ Toxicol Chem 18:2416–2423.

34. Spanggord RJ, Spain JC, Nishino SF, Mortelmans KE. 1991. Biodegradation of 2,4-dinitrotoluene by a Pseudomonas sp. Appl Environ Microbiol 57:3200–3205.

35. Haigler BE, Nishino SF, Spain JC. 1994. Biodegradation of 4-Methyl-5-Nitrocatechol by Pseudomonas sp. Strain DNT. J Bacteriol 176:3433–3437.

36. Nishino SF, Paoli GC. 2000. Aerobic Degradation of Dinitrotoluenes and Pathway for Bacterial Degradation of Aerobic Degradation of Dinitrotoluenes and Pathway for Bacterial Degradation of 2, 6-Dinitrotoluene. Appl Environ Microbiol 66:2139–2147.

37. Akkaya Ö, Pérez-pantoja DR, Calles B, Nikel PI, Lorenzo V de. 2018. The metabolic redox regime of Pseudomonas putida tunes its evolvability towards novel xenobiotic substrates. MBio 9:e01512–18.

38. Pérez-Pantoja D, Nikel PI, Chavarría M, de Lorenzo V. 2013. Endogenous Stress Caused by Faulty Oxidation Reactions Fosters Evolution of 2,4-Dinitrotoluene-Degrading Bacteria. PLoS Genet 9.

39. Ramos JL, Duque E, Maria-Trinidad G, Godoy P. 2002. Mechanisms of solvent tolerance in gram-negative bacteria. Annu Rev Microbiol 56:743–768.

40. Nelson KE, Weinel C, Paulsen IT, Dodson RJ, Hilbert H, Santos VAPM, Fouts DE, Gill SR, Pop M, Holmes M, Brinkac L, Beanan M, Deboy RT, Daugherty S, Kolonay J, Madupu R, Nelson W, White O, Peterson J, Khouri H, Hance I, Lee PC, Holtzapple E, Scanlan D, Tran K, Moazzez A, Utterback T, Rizzo M, Lee K, Kosack D, Moestl D, Wedler H, Lauber J, Stjepandic D, Hoheisel J, Straetz M, Heim S, Kiewitz C, Eisen J, Timmis KN, Düsterhöft A, Tümmler B, Fraser CM. 2002. Complete genome sequence and comparative analysis of the metabolically versatile Pseudomonas putida KT2440. Environ Microbiol 4:799–808.

41. Nikel PI, Chavarría M, Danchin A, de Lorenzo V. 2016. From dirt to industrial applications: Pseudomonas putida as a Synthetic Biology chassis for hosting harsh biochemical reactions. Curr Opin Chem Biol 34:20–29.

42. Yoon JM, Oliver DJ, Shanks J V. 2006. Phytotransformation of 2,4-Dinitrotoluene in Arabidopsis thaliana?: Toxicity, Fate, and Gene Expression Studies in Vitro 1524–1531.

43. Rocheleau S, Kuperman RG, Simini M, Hawari J, Checkai RT, Thiboutot S, Ampleman G, Sunahara GI. 2010. Toxicity of 2, 4-dinitrotoluene to terrestrial plants in natural soils. Sci Total Environ 408:3193–3199.

44. Griest WH, Tyndall RL, Stewart AJ, Caton JE, Vass AA, Ho C-H, Caldwell WM. 1995. Chemical characterization and toxicological testing of windrow composts from explosives-contaminated sediments. Environ Toxicol Chem 14:51–59.

45. Sauer J, Schreiber U, Schmid R, Völker U, Forchhammer K. 2001. Nitrogen starvation-induced chlorosis in Synechococcus PCC 7942. Low-level photosynthesis as a mechanism of long-term survival. Plant Physiol 126:233–43.

46. Korosh TC, Dutcher A, Pfleger BF, McMahon KD. 2018. Inhibition of Cyanobacterial Growth on a Municipal Wastewater Sidestream Is Impacted by Temperature. mSphere 3:1–15.

47. Latifi A, Ruiz M, Zhang CC. 2009. Oxidative stress in cyanobacteria. FEMS Microbiol Rev 33:258–278.

48. Ruiz-güereca DA, Sánchez-saavedra MP. 2016. Growth and phosphorus removal by Synechococcus elongatus co-immobilized in alginate beads with Azospirillum brasilense. J Appl Phycol 1501–1507.

49. Gillet F, Roisin C, Fliniaux MA, Dubreuil AJ, Barbotin JN. 2000. Immobilization of Nicotiana tabacum plant cell suspensions within calcium alginate gel beads for the production of enhanced amounts of scopolin 26:229–234.

50. Bailliez C, Largeau C, Casadevall E, Chimie L De, Physique O, Nationale E. 1985. Applied o. d Microbiology Biotechnology Growth and hydrocarbon production of Botryococcus braunii immobilized in calcium alginate gel 99–105.

51. Srinivasulu B, Adinarayana K, Ellaiah P. 2003. Investigations on Neomycin Production With Immobilized Cells of Streptomyces marinensis Nuv-5 in Calcium Alginate Matrix 4:1–6.

52. Therien JB, Zadvornyy OA, Posewitz MC, Bryant DA, Peters JW. 2014. Growth of Chlamydomonas reinhardtii in acetate-free medium when co-cultured with 1–8.

53. Leino H, Kosourov SN, Saari L, Sivonen K, Tsygankov AA, Aro EM, Allahverdiyeva Y. 2012. Extended H2photoproduction by N2-fixing cyanobacteria immobilized in thin alginate films. Int J Hydrogen Energy 37:151–161.

54. Martínez-García E, Jatsenko T, Kivisaar M, de Lorenzo V. 2015. Freeing Pseudomonas putida KT2440 of its proviral load strengthens endurance to environmental stresses. Environ Microbiol 17:76–90.

55. Löwe H, Sinner P, Kremling A, Pflüger-Grau K. 2018. Engineering sucrose metabolism in Pseudomonas putida highlights the importance of porins. Microb Biotechnol 0:1–10.

56. Löwe H, Schmauder L, Hobmeier K, Kremling A, Pflüger-Grau K. 2017. Metabolic engineering to expand the substrate spectrum of Pseudomonas putida toward sucrose. Microbiologyopen 6:1–9.

57. de las Heras A, Chavarría M, de Lorenzo V. 2011. Association of dnt genes of Burkholderia sp. DNT with the substrate-blind regulator DntR draws the evolutionary itinerary of 2,4-dinitrotoluene biodegradation. Mol Microbiol 82:287–299.

58. Nikel PI, Chavarri M. 2013. Endogenous Stress Caused by Faulty Oxidation Reactions Fosters Evolution of 2, 4-Dinitrotoluene-Degrading Bacteria 9.

59. Romo S, Perez-Martinez C. 1997. The Use of Immobilization in Alginate Beads for Long-Term Storage of Pseudanabaena galeata (Cyanobacteria) in the Laboratory. J Phycol 1073–1076.

60. Wang Z, Ye Z, Zhang M. 2011. Bioremediation of 2,4-dinitrotoluene (2,4-DNT) in immobilized micro-organism biological filter. J Appl Microbiol 1476–1484.

61. Oh S, Seo Y, Ryu K. 2016. Reductive removal of 2,4-dinitrotoluene and 2,4-dichlorophenol with zero-valent iron-included biochar. Bioresour Technol 216:1014–1021.

62. Zavrel T, Sinetova MA, Červený J. 2015. Measurement of Chlorophyll a and Carotenoids Concentration in Cyanobacteria. Bio-Protocol 5:1–5.

63. Shemer B, Yagur-kroll S, Hazan C, Belkin S. 2018. Aerobic Transformation of 2,4-Dinitrotoluene by Escherichia coli and Its Implications for the Detection of Trace Explosives. Appl Environ Microbiol 84:e01729–17.

